# Composite measurements and molecular compressed sensing for highly efficient transcriptomics

**DOI:** 10.1101/091926

**Authors:** Brian Cleary, Le Cong, Eric S. Lander, Aviv Regev

## Abstract

RNA profiling is an excellent phenotype of cellular responses and tissue states, but can be costly to generate at the massive scale required for studies of regulatory circuits, genetic states or perturbation screens. Here, we draw on a series of advances over the last decade in the field of mathematics to establish a rigorous link between biological structure, data compressibility, and efficient data acquisition. We propose that very few random composite measurements – in which gene abundances are combined in a random linear combination – are needed to approximate the high-dimensional similarity between any pair of gene abundance profiles. We then show how finding latent, sparse representations of gene expression data would enable us to “decompress” a small number of random composite measurements and recover high-dimensional gene expression levels that were not measured (unobserved). We present a new algorithm for finding sparse, modular structure, which improves the ability to interpret samples in terms of small numbers of active modules, and show that the modular structure we find is sufficient to recover gene expression profiles from composite measurements (with ~100-fold fewer composite measurements than genes). Moreover, the knowledge that sparse, modular structures exist allows us to recover expression profiles from composite measurements, even without access to any training data. Finally, we present a proof-of-concept experiment for making composite measurements in the laboratory, involving the measurement of linear combinations of RNA abundances. Altogether, our results suggest new compressive modalities in experimental biology that can form a foundation for massive scaling in high-throughput measurements, while also offering new insights into the interpretation of high-dimensional data.

## Introduction

A gene expression profiles is a rich and informative cellular phenotype, which reflects a cell's type, state, and regulatory mechanisms. Beginning with pioneering work on microarray profiling (*1*, *2*) and analysis (*3*–*6*) almost 20 year ago, studies have shown that similarities in gene expression profiles can reveal connections between biological samples, genes co-regulated by shared mechanism, and hierarchical modules of transcriptional programs in diverse organisms, including bacteria (*7*), yeast (*8*), plants (*9*), worm (*10*), fly (*11*), mouse (*12*) and human (*13*).

However, many emerging applications – such as screening for the effect of thousands of genetic perturbations (*14*, *15*), analyzing the genetic basis of variation among individuals (*16*, *17*), large-scale single-cell profiling of complex tissues (*18*, *19*), and diagnostic tests of immune cells in the blood – will require massive numbers of expression profiles, up to hundreds of thousands or more. Efficient means of data acquisition, storage, and computation are thus of critical importance.

A central challenge in gene-expression profiling is the high dimensionality of the data. Mammalian genomes contain approximately 20,000 genes, and mammalian expression profiles are frequently studied as vectors with 20,000 entries corresponding to the abundance of each gene. It is often assumed that studying gene expression profiles requires measuring and analyzing these 20,000 dimensional vectors, but some mathematical results show that it is often possible to study high-dimensional data in low-dimensional space without losing much of the pertinent information. Working in low-dimensional space may offer several advantages with respect to computation, data acquisition and fundamental insights about biological systems.

Image analysis provides an excellent example of the power of low-dimensional representations. In principle, images with 10 Megapixels can be incredibly complex: if the color and intensity of each pixel varies independently, the data require a 10 million-dimensional representation. Yet, the pictures that humans find interesting tend to be highly structured and, as a result, highly compressible. Pictures can be expressed in terms of their Fourier Transform (or the closely related discrete cosine transform) (*20*), which often leads to a much more compact representation because most Fourier coefficients are small enough to ignore. Wavelets provide even better representations (*21*), because they can efficiently capture features corresponding to edges and fields. As a result, the high-dimensional representation of an image can often be reconstructed from a low-dimensional embedding in a space of wavelet-based features. These representations are the basis for compression algorithms, such as JPEG. Overall, dimensionality reduction works in image analysis because a limited ‘dictionary’ of functions does a good job of capturing most of the information that is relevant to human cognition in the images of interest.

Approaches that employ dimensionality reduction are already applied in some contexts to study gene expression, including for data interpretation (*22*–*25*) and algorithmic efficiency (*26*–*28*). Methods of bi-clustering (*6*, *25*) and gene signature analysis (*29*) have identified sets of genes (often termed “modules”) with coordinated expression levels in a subset of conditions. Membership in modules helps biologists annotate genes of unknown function (through “guilt by association”) (*30*), identify functional processes affected in different physiological conditions (*29*, *31*) and disease, and infer regulatory mechanisms (*29*). In some cases, a much more limited subset (“signature”) of genes can be identified, which, when measured in new samples, can be used together with the earlier profiling data to estimate the abundance of the remaining unmeasured (unobserved) genes in these new samples (*32*–*36*). Most recently, with the advent of massively parallel single-cell RNA-Seq, it is becoming increasingly common to use shallow RNA-Seq data of each sample (here, a single cell) to draw inferences about the complete gene expression profile (*18*, *37*–*41*). It has been shown that analyzing tens of thousands of cells at a low depth (and cost) per cell still allows for the recovery of meaningful biological distinctions on cell type (*18*, *39*, *41*) and state (*37*, *38*, *40*), and that this is attributable to inherent low-dimensional structure in gene expression data (*42*).

Yet, current methods do not capture the full power of dimensionality reduction. For example, bi-clustering and signature analysis provide biological understanding, but these semi-categorical methods, which produce sets or ranked lists of genes, don't directly suggest any means of compressive data acquisition, since fully quantitative descriptions are required to convert between compressed and decompressed data. On the other hand, methods such as Principle Component Analysis (PCA) provide quantitative descriptions, but, as we show below, the reduced dimensions are often limited in their biological interpretability.

Current approaches also have important limitations with respect to experimental design. Approaches that aim to identify small signatures to be used for subsequent experimental measurements tend to be limited by nature of the training data, the model used for imputation, and the quality of their imputation. Approaches that rely on shallow sequencing tend to perform poorly at capturing information about low to moderately expressed genes, including key genes like transcription factors (*42*).

Here, we seek to establish a rigorous link between biological structure, data compressibility, and efficient data acquisition. We draw on a series of advances over the last decade in the field of mathematics—specifically, the theory of compressed sensing (*43*, *44*) (reviewed in (*45*)), which formalizes a framework for designing low-dimensional (compressed) measurements and using them to recover a structured, high-dimensional signal (here, a gene expression profile with latent modular structure). Interestingly, the data compression can often be performed simply by using random projections into low-dimensional space.

Our main results are as follows: (*1*) We show that it is possible to quantify the similarity between samples even after projecting the 20,000-dimensional gene-expression data into as few as 100 dimensions, each a random combination of gene expression levels. (*2*) We propose that gene-expression data itself might be collected directly in these 100 dimensions, by performing “composite measurements” that involve measuring linear combinations of genes. (*3*) We show that high-dimensional gene-expression data has a sparse, modular structure. (*4*) We present a new algorithm for finding such modular structure, which improves the ability to interpret gene expression profiles in terms of small numbers of active modules. (*5*) We show that the modular structure is sufficient to recover gene expression profiles from composite measurements (with ~100-fold fewer composite measurements than genes). (*6*) We show that the knowledge that sparse, modular structures exist allows us to recover expression profiles from composite measurements “blindly”, that is even without access to any training data. (*7*) We present a proof-of-concept experiment for making composite measurements in the laboratory, involving the measurement of linear combinations of RNA abundances.

Overall, the results suggest new approaches for both experiments and interpretation in genomics and biology.

## Expression Data

In our analyses below, we will use 40 publically available datasets containing a total of 24,374 unique expression profiles (**table S1**). Thirty-six of the datasets are from bulk tissue samples: the GTEx collection of human tissues (8,555 profiles (*16*)); the ImmGen dataset of mouse hematopoietic and immune cell types (214 profiles (*46*)), and The Cancer Genome Atlas (TCGA, 33 datasets from 33 cancer types analyzed separately, as well as a “combined” TCGA dataset containing all 10,554 profiles from all cancer types (*47*)). The remaining four datasets are from single-cell mRNA expression profiles (scRNA-Seq); they consist of datasets from studies of mouse cerebral cortex (3,005 cells) (*48*); adult mouse primary visual cortex (1,809 cells) (*49*); intestinal epithelial cells (192 cells) (*50*); and a rare population of human radial glial cells (45 cells) (*51*).

While these data were generated with diverse technologies (bulk RNA-Seq, scRNA-Seq, and microarrays), we performed only minimal normalization to ensure robustness to the method of data collection (SOM). Specifically, the only normalization was to put a ceiling on abundance at the 99.5^th^ percentile in each dataset to avoid performance statistics that are skewed by few genes with extremely high expression.

## Similarity of expression profiles: Theory

We start with a simple question: How can we efficiently compare expression profiles? Suppose that we have expression profiles from a large collection of *n* samples, represented as points in 20,000-dimensional space (where we let the number of genes, *g* = 20,000). We can quantify the similarity between a pair of samples by calculating the Euclidean distance between the corresponding pair of points. (Euclidean distance is equivalent to correlation if the samples have been suitably normalized.) To do this for all pairs, the total computational “cost” would be proportional to:

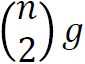

An elegant mathematical result of Johnson and Lindenstrauss (*52*), however, suggests an alternative approach, which incurs a far lower computational cost. The result, along with subsequent proofs (*53*, *54*), states that the distances among *n* points in a high-dimensional space can be well preserved with very high probability if the data are projected onto a randomly defined *m*-dimensional subspace, where *m* is on the order of log *n*.

In our case, projecting a point in 20,000-dimensional gene-expression space onto a *single* dimension amounts to taking a linear combination (weighted sum) of the 20,000 gene-expression levels. In a *random* projection, the 20,000 weights in the linear combination are chosen randomly. (Specifically, the weights are chosen as identically and independently distributed Gaussian random variables with mean 0 and standard deviation 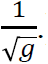 Projecting onto multiple dimensions simply means taking multiple such linear combinations (**Fig. 1A**). We write this formally as:

**Figure 1:**
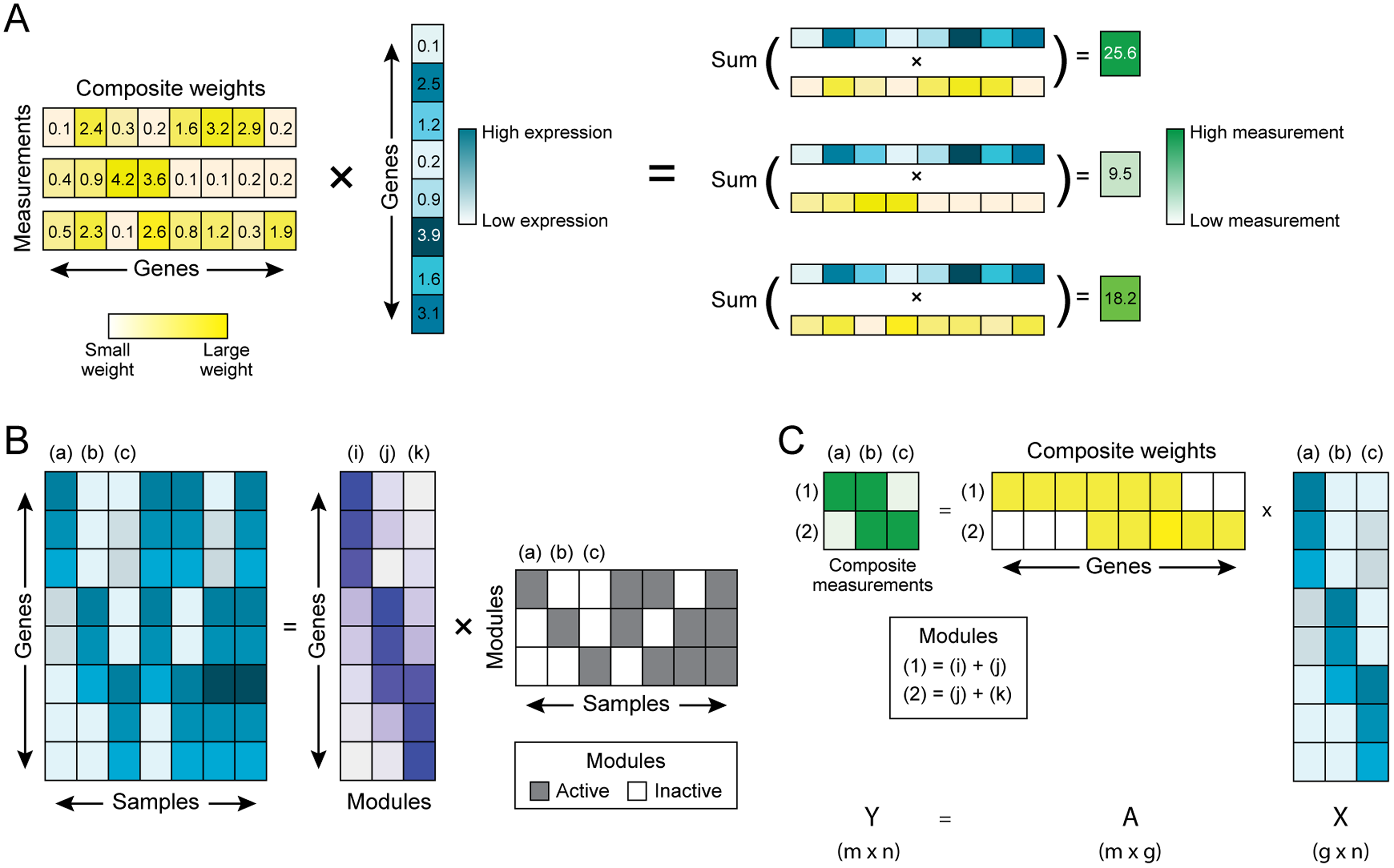
Composite measurements of sparse module activity. (A) Composite measurements are “composed” from the abundance levels of multiple genes. Schematic example of three composite measurements (green, right) constructed from one vector of gene abundances (cyan). Each measurement is a linear combination of gene abundances, with varying weights (yellow) for each gene in each measurement. (**B**) Decomposition of gene abundance across multiple samples by the activity of gene modules. The expression of genes (rows) across samples (columns) (left cyan matrix) can be decomposed into gene modules (purple matrix; rows: genes; columns: modules) by the modules' activity (grey matrix, rows) across the samples (grey matrix; columns). If only one module is active in any sample (as in samples a, b, and c) then two composite measurements are sufficient to determine the gene expression levels (part **C**). One such measurement (1) is composed from the sum of modules (i) and (j), and another (2) is composed from the sum of modules (j) and (k).

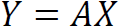

where the matrix *X* (*g* × *n*) represents the original gene-expression values of *n* samples in *g*=20,000-dimensional space; the matrix *A* (*m* × *g*) represents the weights of *m* random linear combinations; and the matrix *Y* (*m* × *n*) represents the samples in low-dimensional space. (Note: in order for distances in *Y* to be on the same scale as those in *X* we need to multiply by a constant factor. However, since below we will be concerned with the *correlation* – which adjusts for scale – between pairwise distances in *Y* and *X*, we ignore this scaling without loss of generality.)

The computational cost of (1) projecting the data into *m*-dimensional space and (2) calculating Euclidean distances between the points in *m*-dimensional space is

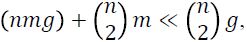

since *m* is of order log *n*.

In addition to providing an efficient way to *analyze* data, the mathematical approach suggests the possibility of an efficient way to *collect* data: rather than measuring the expression level of each of 20,000 genes individually, perhaps we could measure *m* linear *combinations* of genes—which we will refer to as “composite measurements.” If composite measurements had the same cost as individual gene measurements, there might be considerable cost savings in data collection, and we could use the low-dimensional data to learn about the correlations among the expression profiles of the samples. (Moreover, as discussed below, we can use low-dimensional data to recover information about the abundance of the 20,000 individual genes.) We return to this idea in the final portion of the Results section, where we present some pilot experiments addressing whether such a scheme might be technically feasible.

## Similarity of expression profiles: Application

We set out to test the method by applying it to each of our 40 datasets. We compared (1) the pairwise distances between samples as represented in the low-dimensional embedding (compressed dataset), *Y*, with (2) the pairwise distances in the original high-dimensional space (full original dataset), *X*. With only *m*=100 random composite measurements, the Pearson correlation coefficient (averaged across the 40 datasets) between the distances in the low-and high-dimensional spaces was 94.4%. With *m*=400 random measurements, the average correlation was 98.5%.

In a subsequent section, we will discuss how linear combinations might be experimentally collected in the laboratory. To allow for this possibility, we will assume the addition of some noise. Specifically, we imagine that we have a noisy estimate of each gene in each sample, that the noise process for each gene in each sample is identically and independently distributed (i.i.d.) as a Gaussian, and that the overall signal-to-noise ratio (SNR) is 2. Thus, we write noisy composite measurements as:

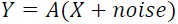

With *m*=100 noisy measurements, the average correlation between low and high dimensional distances was 84.5%. With 400 noisy measurements, the average correlation was 95.1%. To allow for the possibility of data collected by means of noisy composite measurements, we include the addition of noise (SNR=2) in all calculations below. (The results below are robust to other choices of SNR, data not shown.)

In addition to finding that pairwise distances are preserved, we find that the clustering of samples in the low-dimensional space approximates the clusters found in the high-dimensional space. For example, the GTEx dataset consists of 8,555 samples from 30 different tissues. When we generate 30 clusters in high-dimensional space, we find that samples are grouped by tissue and that this pattern is mirrored exceptionally well in the low-dimensional data from the 100 random projections (**Fig. 2**). For example, the 412 heart samples are members of a single cluster in the high-dimensional data, whereas 411 of these samples lie in a common cluster in the low-dimensional data. For those tissues split into multiple clusters in high dimension, we find that they are split into a similar number of clusters, with similar groupings, in low dimension. For example, for the colon samples, the effective numbers of clusters (calculated using Shannon Diversity) was on average 3.32 clusters in high dimension, and 2.88 clusters in low dimension. Using all of the clusters, we can quantify the similarity between the low-and high-dimensional results in terms of the “mutual information”—that is, the reduction in uncertainty about the high-dimensional clustering, given the low-dimensional clustering. In the GTEx data, the uncertainty about a pair of samples falling in the same cluster in high dimension is reduced by 87% if we are given the clusters found in the 100-dimensional data.

**Figure 2:**
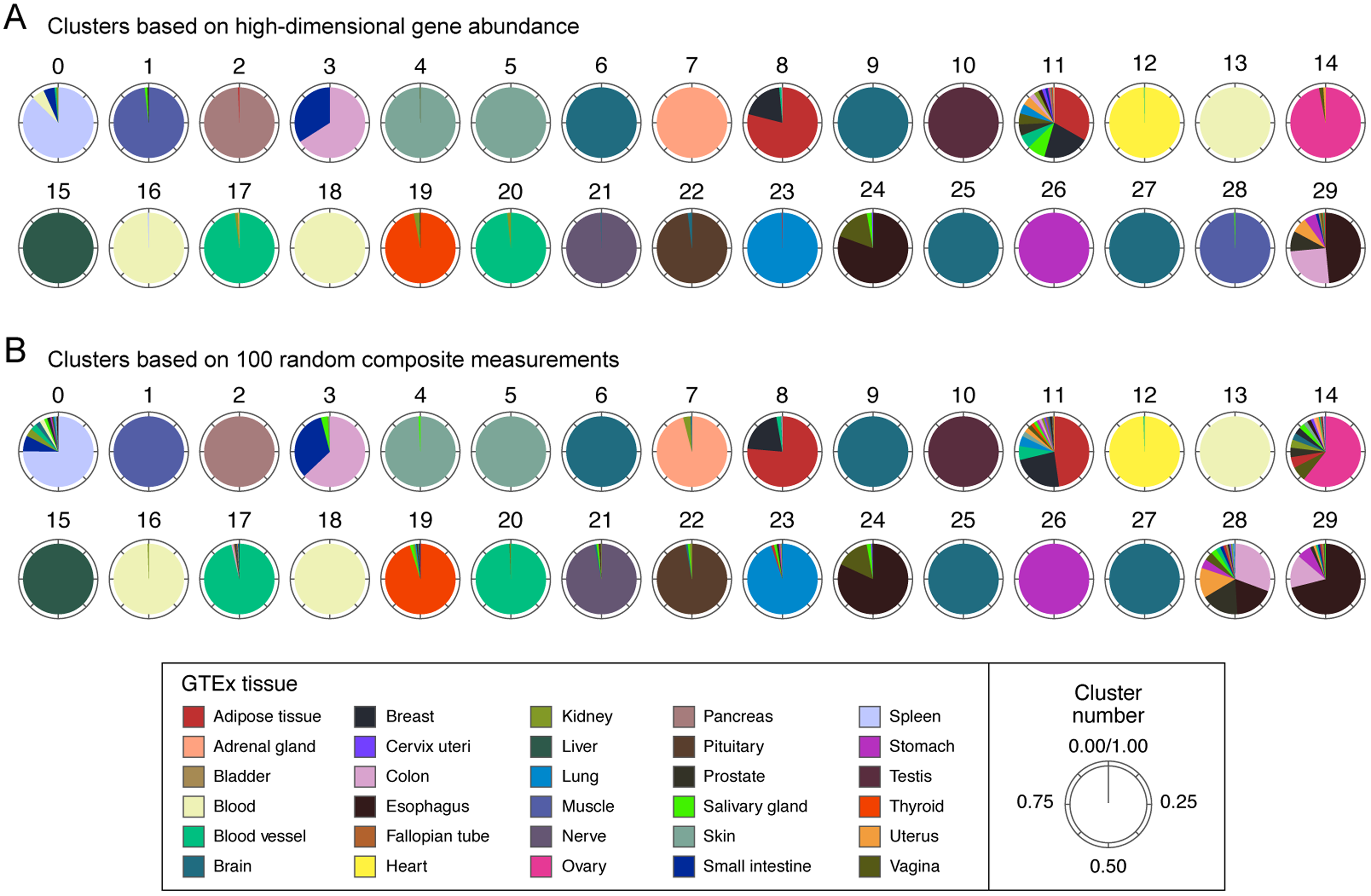
Comparable clustering of GTEx samples by high-and low-dimensional measurements. GTEx samples were clustered into 30 groups (0-29) on the basis of 14,202 gene expression levels (**A**), or 100 random composite measurements (**B**). Each pie chart corresponds to one cluster. Colors correspond to individual tissues (legend), and the area covered by each color in a pie depicts the fraction of samples in a cluster that were derived from a given tissue.

In some datasets, clusters are not as easily distinguished as in the GTEx dataset, particularly where the samples are all derived from the same tissue. For example, within each of the 33 individual TCGA tumor types, the average reduction in uncertainty was 36%. (Across all TCGA tumor types together, the reduction was 47%.) This reflects some loss of information about clustering in low dimension, but it is also partially due to the fact that the high dimensional clusters *themselves* are not inherently well-separated (for example, high-dimensional clusters within TCGA tumor types change more dramatically after the addition of noise; **fig. S1**). Even so, in these cases, the distances themselves are well preserved from high to low dimension (84% correlation).

## The role of modular structure in recovering gene abundances from low-dimensional data: Theory

Next, we consider a harder problem: can we recover the abundance of each of the 20,000 genes from random composite measurements? In the field of compressed sensing, it has been shown that this is possible provided that the high dimensional data possess a latent sparse structure (*43*, *44*). Encouragingly, as we discussed, extensive studies of gene expression programs suggest that they are highly structured. In this section, we will begin by reviewing the challenges posed by this problem and the need to identify latent structure in order to find a solution. We will then clarify our definition of modular structures and discuss several algorithms for finding these. After empirically exploring the inherent structure of gene expression datasets in the next section, we will return to developing a theoretical framework for the recovery of gene expression profiles from composite measurements using compressed sensing.

The challenge in recovering high-dimensional data (expression profiles) from low-dimensional observations (composite measurements) centers on the underdetermined system of equations:

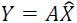

where *Y* and *A* are known, and we wish to estimate *X̂*, the predicted gene expression levels in 20,000-dimensional space. If gene expression is arbitrarily complex (that is, if the abundance levels of each gene may vary across samples independently from all other genes), it is impossible to infer the gene expression pattern from a small number of measurements. On the other hand, if some genes are co-regulated and thus their abundances co-vary, then, intuitively, the problem should be easier. In an extreme case, if two genes are perfectly correlated, we need only measure one in any given sample. More generally, if gene expression is organized into co-varying modules, we can consider the alternative – and potentially easier – problem of using composite observations to determine the activity level of each module in each sample. The more that gene expression follows a ‘modular’ structure, the better the prospects of recovering the 20,000 abundances from far fewer measurements.

As a concrete example of what the hidden modular structure of gene expression might look like, we might imagine that each cell can choose from a collection of 500 fundamental gene expression programs and may activate at most 15 of them at a time. Adopting the terminology of compressed sensing, we would refer to the 500 gene expression programs as the “dictionary” and would say that cells used gene expression patterns that were “*k*-sparse”, with *k* = 15. (For practical purposes, it will be enough that cells' gene expression patterns can be approximated by such a representation — that is, they are *approximately* k-sparse.)

If expression were truly (or approximately) structured in this way, this could allow us both to (1) better design ways to compress biological measurements, and (2) better understand the organization and regulation of gene expression and biological processes.

With respect to the first point above, consider the following toy example of inferring modular activity from “under-sampled” measurements. Suppose that the dictionary has only three fundamental programs, and each cell activates only one of these programs (**Fig. 1B**). With only two measurements — one composed of the sum of programs A and B, the other of the sum of programs B and C—we could infer which of the three modules is active (**Fig. 1C**). Because the dictionary is both small and sparsely used, we can recover the gene expression from a number of measurements that is smaller than both the number of genes and the number of modules (that is, 2 *v*s. 20,000 and 3). Current approaches to addressing such sampling challenges often involve using shallow sequencing or a set of signature genes (*32*, *33*, *35*, *55*). Below, we will show that we can improve both conceptually and practically upon these methods with a distinct mathematical framework.

With respect to the second—and biologically more fundamental—point above, knowledge of a dictionary of possible expression programs and their activation across cell types could help identify the key functional modules active in biological samples and help infer the regulatory mechanisms that control them.

To address both goals, we begin by formalizing the notion of modular activity in the following way: if *x_t_* denotes the vector of 20,000 gene-expression levels in sample *i*, we want to write these levels as a weighted sum of the 500 hypothetical modules:

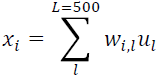

where *u_t_* denotes the weights of 20,000 genes in module *l* and the coefficients *w_i,l_* define the activity level of module *l* in sample *i*. We thus seek an algorithm to take the matrix *X* of observed expression levels and express it as the product of a dictionary *U* of expression modules and a matrix *W* of module activity levels. Posed in this way, this becomes a problem of matrix factorization: *X* = *UW* (**Fig. 3A**).

**Figure 3:**
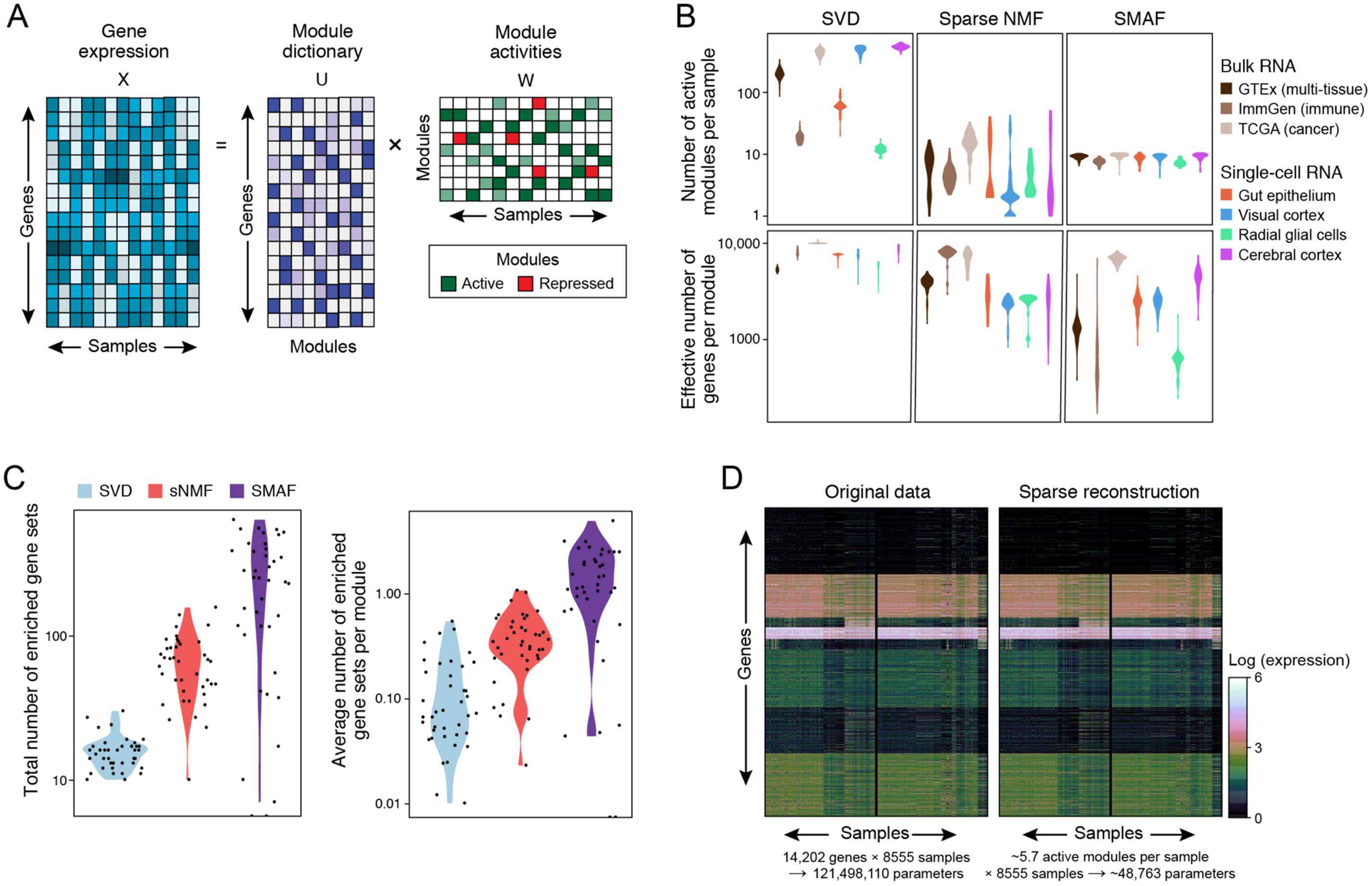
Sparse Modular Activity Factorization (SMAF) for gene expression. (**A**) Decomposition with a module dictionary. A gene expression matrix, × (left; rows: genes; columns: samples), is decomposed into a module dictionary, U (middle; rows: genes; columns: modules), and module activity levels in each sample, W (right; rows: modules; columns: samples). (**B, C**) Performance of different matrix decomposition algorithms. (**B**) Violin plots of the distribution of the number of active modules per sample (top), and the effective number of genes per module (bottom) for each of three methods, across different datasets (x axis). (**C**) Violin plots of the distribution of the total number of enriched gene sets across all modules within a dataset (left), and the average number of enriched gene sets per module (right), for each of the three different algorithms. Each dot represents one dataset. (**D**) Reconstructed high-dimensional gene expression levels. Heat maps show, for the GTEx dataset the original gene expression profiles (left) and the profiles reconstructed from the sparse module activity levels from SMAF.

There are several methods of matrix factorization that are commonly used for gene expression analysis. Two of the best-known algorithms are Singular Value Decomposition (SVD) (*22*) and nonnegative matrix factorization (NMF) (*24*). For the purpose of compression, we desire an algorithm that can accurately represent gene expression with a small number of active modules (*i.e*. very few nonzero *w_i,l_* coefficients). The general versions of SVD and NMF are not guaranteed to accomplish this, but modified versions, such as sparse NMF (*56*), incorporate sparsity constraints to enforce such behavior.

To reflect our current understanding of the functional and regulatory underpinnings of gene modules and to increase the biological interpretability of the resulting dictionary, we define four desirable features: (**1**) Sparse usage of modules: as discussed above; (**2**) Restricted modularity: the number of genes in any module should be relatively small, and, correspondingly, genes should not participate in too many modules; (**3**) Biological coherence: different modules should represent distinct pathways or programs, and should not overlap too much; and (**4**) Compactness: the total number of modules should not be too large. This list provides criteria for evaluating the results of different algorithms for finding modules and modular activity. We first evaluate the established SVD and sparse NMF algorithms, and use these results to motivate the development of a new algorithm that is specifically tailored to our criteria.

## Modular activity in gene expression profiles: Results

To assess the performance of the SVD and sparse NMF algorithms by our criteria, we used each method to calculate module dictionaries and module activity levels for each of the 40 datasets (SOM). The SVD algorithm places no constraint on sparsity, while sparse NMF (sNMF) uses a “soft” constraint that results in a generally sparse activity matrix, albeit without an explicit cap of, *k* the number of non-zero modules per sample.

With SVD, we found dictionaries with an average of 312 modules; they fit the data well, with a 99.2% fit on average across all datasets (**table S2**). As expected, the representations were not sparse: samples were frequently represented by a linear combination of hundreds of modules.

With sNMF, we found an average of 326 modules. The results not only fit the data well (88.6% fit on average across all datasets, **table S2**), but produced sparse solutions, with an average of 5.1 active modules per sample (**Fig. 3B, fig. S2A**).

Neither method, however, produced restricted modularity: in both cases, most modules involved *thousands* of genes. To quantify this we define the total module ‘weight’ as:

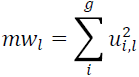

(*i.e.*, as the squared sum of coefficients in a module). Considering the genes with largest (absolute value) coefficients, with SVD we needed 7,417 genes on average to capture 99% of the module ‘weight’, and with sNMF we needed 6,564 genes (**Fig, 3B, fig. S2B**). In addition, each gene was represented in *hundreds* of modules (SVD: 224 modules per gene on average; sparse NMF: 215 modules per gene).

To assess biological coherence, we tested the modules for enrichment in functional gene sets (*57*, *58*) (SOM). Focusing on the five most enriched sets for each module (with FDR q-value < 0.05), we found that these enriched gene sets largely overlap between modules; we define the number of ‘unique’ enriched gene sets in a dictionary to be the set of terms enriched in at least one module, and the number of unique gene sets per module as the total number of unique enrichments in a dictionary divided by the total number of modules. We find only 0.13 and 0.41 unique gene sets per module, in SVD and sNMF, respectively (**Fig. 3C**). If we truncate the list of genes for each module by considering only those genes comprising 50% of the total ‘weight’ (as opposed to 99% above), the number of unique sets per module increases (to 1.04 and 1.26, respectively), but these truncated modules do a significantly worse job of quantitatively accounting for the original data (for instance, in GTEx, the fit is reduced from 99% to 49%).

From these results, it is unclear whether the failure to obtain modules with the features of restricted modularity and biological coherence is due to limitations of computational methods or inherent properties of the biological data. Because earlier studies, especially in yeast (*8*, *30*) but also in human (*13*, *29*, *31*) have found coherent modules, we hypothesized that an algorithm devised with these considerations in mind could perform better.

We therefore developed a new algorithm, Sparse Module Activity Factorization (SMAF), with the aim of better achieving the four criteria listed in above (SOM). SMAF has two important properties. First, SMAF requires both the module dictionary and the module activity levels to be sparse—with a soft constraint on sparsity in the dictionary (*i.e.* a penalty for including more nonzero terms – see SOM) and a hard constraint on *k*-sparsity in the activity levels (*i.e.* at most *k* nonzero terms per sample (column of W)). Second, SMAF requires that each entry in the module dictionary consists of nonnegative coefficients (*i.e. u_i,l_* ≥ 0 *for all i*, *l*)—because these values are meant to be interpreted as “membership” in a module. However, the activity levels of each module, determined by coefficients *w_i,l_* for module *l* in sample *j*, may be either positive or negative— reflecting the notion that modules can either be activated or repressed. With at most *k* (active or repressed) modules contributing to each sample, the observed expression levels would reflect a linear combination of sparse modular activity.

We applied SMAF to each of the 40 data sets, testing its ability to create dictionaries that provide a good 10-sparse (*k*= 10) fit to the expression data. SMAF yielded a good fit (average of 93%, **table S2**), while producing sparse dictionaries (average of 248 modules per dictionary) of modules that are sparsely active and biologically coherent (average of 1.45 uniquely enriched gene sets per module without truncation); and, consequently, a much greater number of gene sets enriched in at least one module (average of 372 total unique enriched gene sets per dataset versus 15 with SVD and 63 with sparse NMF) (**Fig. 3B-D**). Using a truncated list of genes (comprising 50% of the weight) for each module further improved the enrichments, but only moderately so (from 1.45 unique sets per module to 1.62). SMAF's excellent performance in fitting the data supports the hypothesis that sparse structures can provide a good description of gene expression data.

For example, consider a sample (TCGA-A8-A09R-01) from the TCGA-Breast invasive carcinoma (TCGA-BRCA) dataset. The 10-sparse SMAF reconstructed profile of this sample had a 92.6% correlation with the original values. Based on the most significant gene set enrichments, the modules represent: (**1**) a set of genes involved in telomeric maintenance and the genes within a chromosomal region (chr17q21) with aberrant overexpression in this tumor; (**2**) a set of genes involved in the maintenance of chromosome structure and genes that are specifically upregulated in ESR1 positive breast cancers (this sample is ER+); (**3,4**) the genes within several chromosomal regions with aberrant overexpression – these partially overlap with module (*1*) (*e.g*., chr17q21-q25), but include additional regions as well (chr17q11 and chr18q21(note that the TCGA database indicates up-regulated gene expression in this tumor along chr17q11-q25, although genetic analysis by TCGA does not report a large-scale copy number amplification in the region); (**5**) genes that direct the differentiation of mammary stem cells; (**6**) genes involved in the breakdown of an extracellular matrix; (**7**) genes involved in leukocyte activation and in cell motility (likely reflecting immune infiltrate in this bulk tumor sample); (**8**) genes within an amplified chromosomal region in this tumor (chr8p11); and (**9**) genes that respond to stimulus by type I interferons (reflecting either intrinsic malignant cell responses or additional immune infiltration). The one repressed module consists of (**10**) genes involved in DNA damage repair (which can lead to chromosomal instability);

As a second example, we picked a random sample from the GTEx dataset (GTEX-1399T-2426-SM-5L3FJ), derived from skeletal muscle. The expression levels calculated from a linear combination of the 10-sparse SMAF profile had a 98.72% correlation with the original values. Based on gene set enrichment, the modules are associated with: (**1**) striated muscle contraction; (**2**) formation of myofibrils and progression of muscle structure; (**3**) a module of housekeeping genes that is broadly expressed in many tissues; (**4,5,6**) three modules of housekeeping genes that are specifically active in muscle samples (the genes are broadly expressed and participate in processes such as transcription and translation, but the relative abundances have muscle specific profiles); (**7,8**) repression of a respiratory electron transport chain module that is specific to a subset of muscle samples, and activation of a similar module that is specific to a different subset (including this sample); (**9**) genes involved in muscle sliding and ATPase activity; and, (**10**) repression of genes specifically involved in cardiac muscle contraction.

Finally, we randomly chose a single-cell profile from adult mouse cortex (Vip_tdTpositive_cell_15) (*49*). The 10-sparse SMAF profile showed 99.3% correlation with the original expression data. The modules are associated with: (**1**) active transport of protons against a gradient; (**2**) metabolism and the release of energy; (**3**) repression of a subset of genes that establish an electrochemical gradient; (**4**) housekeeping genes, specifically including those involved in the targeting of proteins to a membrane; (**5,6**) two more modules associated with the establishment of a gradient; (**7,8**) two modules involved in cell-to-cell communication; (**9**) repression of genes involved in the progression of neural structures through development; and, (**10**) ion transport across a membrane.

Taken together, we found that the gene expression profiles examined, like images, can be well approximately by a relatively small number of features. The list of features (*i.e.*, the module dictionary) used across all samples may be relatively large (in the 100's or 1000's), but the expression levels in any one sample can be reasonably well explained by the activity of a small number of (active or repressed) modules. Furthermore, an algorithm (SMAF) constrained by assumptions of sparsity and modularity can capture most of the information in gene expression and the discovered dictionary of gene modules appears to be biologically meaningful. In contrast, the top modules for these samples in SVD and sNMF are redundant and more difficult to interpret (**table S3**).

## Using sparse modularity to recover gene expression profiles with compressed sensing: Theory

We now return to the question of whether we can recover high-dimensional gene expression profiles from low-dimensional data. Suppose that (**1**) we already know a high-dimensional dictionary *U* (or learn it from initial training data), and (**2**) we have a set of low-dimensional composite measurements of gene expression, *Y* = *A*(*X* + *noise*) produced with a known random matrix *A*. We ask whether it is possible to use this information to learn the weighted activity levels *W* for the modules corresponding to the high-dimensional data—and thereby recover the high-dimensional gene expression data *X* ≈ *UW* (**Fig. 4A,B**). In computer science, this is referred to as *compressed sensing*. (In a later section, we address the even harder problem of recovering *X* without information about *U*—referred to as *blind* compressed sensing.)

**Figure 4:**
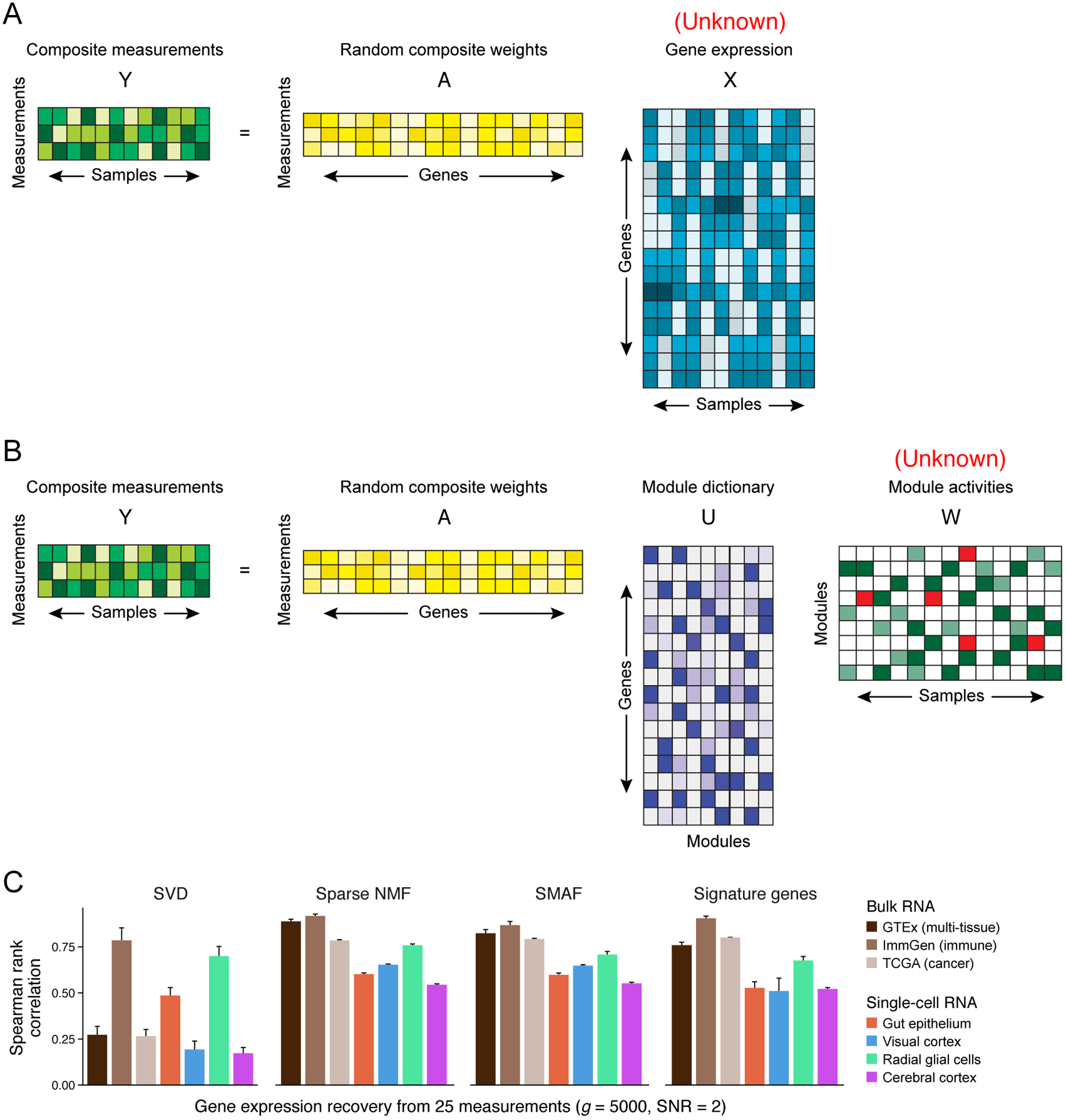
Compressed sensing of gene module activity levels. (A) With a small number of composite measurements, Y, gene expression levels, X, cannot be directly inferred using only information about the compositional weights, A. (B) If gene expression can be decomposed into a module dictionary, U, that is sparsely activated in any sample, then a small number of composite measurements may be sufficient to determine these activity levels. **(C)** Performance of module dictionaries from three different algorithms (SVD, Sparse NMF, and SMAF) in compressed sensing. Each algorithm was used to find module dictionaries in training data. Then, in testing data, 25 simulated noisy composite measurements of 5,000 genes were used to estimate module activity levels of each sample in a dataset. These values were then used to predict the 5,000 expression levels. Bar plot shows for each module dictionary the Spearman rank correlation coefficient (Y axis) between predicted and actual levels, and error bars represent one standard deviation across 50 random trials. These results were compared with simulations of 25 signature gene measurements, which were used to predict the levels of the remaining genes based on models built in training data.

More formally, our process is as follows (SOM). We divide each data set *X* into a set of training samples, *X_training_*, consisting of 5% of the data, and a test set, *X_training_*, consisting of 95% of the data. Using the training set, *X_training_*, we use each of the three algorithms (SVD, sNMF and SMAF) to calculate a module dictionary via matrix factorization:

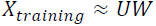

We simulate random compositional measurements on the test samples:

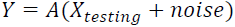

where the matrix *A* defines the random composition of *g* genes in each of *m* measurements (as before, with *m* ≪ *g*). We analyzed a randomly selected set of 5,000 genes (rather than 20,000, in order to reduce the running time across a large number of random trials) and applied noise with signal-to-noise ratio of 2.

For a module dictionary (*U_SVD_*, *U_NMP_*, or *U_SMAP_*), we seek to find the module activities, *Ŵ*, that best fit our composite observations, such that (**Fig. 4B**):

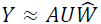

This optimization is performed while enforcing *k*-sparsity in *Ŵ*, so that there are no more than 15 active modules per sample. (Empirically, allowing slightly larger *k*—15 here as opposed to 10 when learning the initial module dictionary from the training data—improves the fit when training modules may not be perfectly representative of testing samples). Finally, we use the module activity coefficients in each testing sample to compute predicted gene expression values, *X̂* = *UŴ*, and compare these predicted values with the actual values. The results are three sets of predictions, *X̂_SVD_*, *X̂_NMP_*, and *X̂_SMAP_*.

## Compressed sensing: Results

We applied this approach to each of the 40 datasets. Successful recovery of gene expression from compressed measurements rests on samples actually having a sparse representation in the given dictionary—that is, on there being not too many active modules in each sample. As predicted by our earlier analysis, this approach indeed worked well for dictionaries based on sNMF or SMAF, but not SVD (**Fig. 4C, fig. S3**). For example, in the GTEx data set with 25 random composite measurements, the results of compressed sensing showed an average Spearman rank correlation with the original values (calculated across all genes and testing samples) of 26% for SVD, 88% for sNMF and 82% for SMAF (**Fig. 4C**). (In fact, with only 10 composite measurements we obtained surprisingly good accuracies of 83% for sNMF and 79% for SMAF (**fig. S4**).)

Some features of gene expression variation were not captured as well as others. Most notably, on average, the approach more accurately estimated the relative abundance of genes within an individual sample than the relative abundance of an individual gene across many samples. For example, in the GTEx data with 25 composite measurements the “overall” correlation across all genes and samples was 82%, while the average correlations within and across samples were 85% and 47%, respectively (**fig. S5**).

The performance was also generally worse for expression datasets from single-cells than from bulk tissue (average Spearman correlation of 62% vs. 87%, for 25 composite measurements). This can be partially explained by the effect of single cell “zero-inflation” (*i.e.*, the absence of sequencing reads for expressed genes) (*59*) and the skewed distributions in single cell expression profiles: in these data the observed abundances are generated by something close to a Poisson process—with many zeros, but also some very large counts—but our optimization methods effectively model the data as normally distributed. When the expression data are highly non-normal we expect a larger difference between the Pearson and Spearman correlation statistics (Pearson implicitly assumes normality, and may be inflated relative to Spearman when the data are non-normal). Indeed, the difference between Spearman and Pearson correlation is most drastic in single cell datasets (**fig. S6**). In addition, there are two sources of noise that could affect performance, regardless of the choice of statistic. First, single-cell RNA Seq may capture intrinsic noise (*e.g.*, transcriptional bursting (*60*)), which is not well captured by modules shared across cells. Second, current single-cell RNA-Seq methods likely have greater technical variability than bulk RNA methods, which similarly would create cell-intrinsic noise that may not be well captured with shared gene modules. The experimental method that we discuss below has been designed to address several sources of technical variability that are particularly significant in single cell RNA-Seq.

Despite these limitations, the results above demonstrate that a small number of random composite measurements are sufficient to provide a good approximation of the abundance of thousands of genes (in this case, 25 measurements for 5,000 genes).

## Recovering gene modules and activity levels from random compositional measurements without knowledge of expression patterns: Theory

In the previous section we saw that it is possible to recover high-dimensional gene expression levels from low-dimensional composite measurements—provided that we are given a high-dimensional dictionary *U*, or at least a training set of high-dimensional data from which we can learn *U*. Now we ask: Suppose we have neither of these things. Can we learn a dictionary *U* from only the low-dimensional composite measurements? In other words, in the equation:

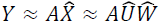

can we learn the module dictionary *Û* based only on knowledge of the low-dimensional data *Y* and projection matrix *A*? (Once we have the dictionary *Û*, we can apply the methods of compressed sensing described above to recover the module activity levels, *Ŵ*, and the original gene abundances, *X̂* (**Fig. 5A**).)

**Figure 5:**
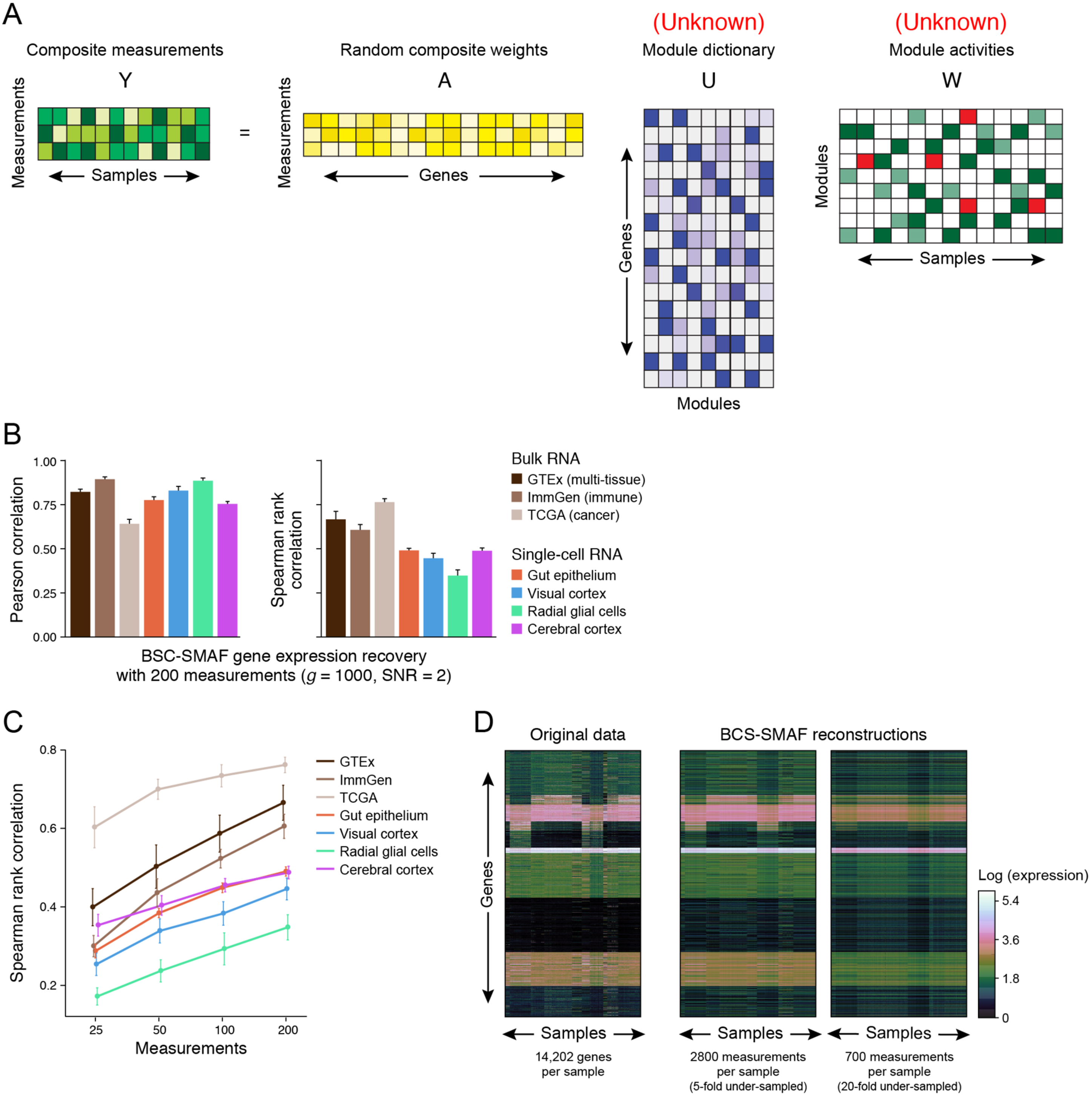
Blind compressed sensing (BCS) of gene modules. (A) In BCS, composite measurements are used to learn both the module dictionary and the module activity levels. (B-D) Performance of BCS. For each dataset, 200 noisy composite measurements of 1,000 genes were simulated. BCS-SMAF was used to learn the dictionary and activity levels, and to predict the 1,000 gene abundances. (B) Bar plots show the Pearson (left) and Spearman (right) correlation coefficients (Y axis) between predicted and actual gene abundances, and error bars represent one standard deviation across 50 random trials. (**C**) Spearman correlation coefficients (Y axis), as in (B), but for varying numbers of composite measurements (X axis). (**D**) Original (left) expression levels for all 14,202 genes in GTEx and their corresponding predictions by applying BCS-SMAF to 2,800 (middle) and 700 (right) composite measurements.

In the compressed sensing field, this problem is called “blind compressed sensing” (BCS). Remarkably, the problem can be solved under certain conditions (*61*, *62*). Specifically, (**1**) the module activity levels must be *k*-sparse; (**2**) we need access to a large number of low-dimensional observations, *Y*; and (**3**) the dictionary itself cannot be too complex. (In blind compressed sensing, various criteria are used to assess “complexity.” Here, we will define complexity in terms on the total number of non-zero coefficients used across the entire dictionary.)

To learn a dictionary, we can proceed as follows:

1. Cluster samples based on low-dimensional observations. As shown above (**Fig. 2**), these clusters should be good approximations to clusters that could be found from high-dimensional data.
2. For every sample, make a crude approximation of the high-dimensional data by taking the pseudo-inverse: *X̂_initial_* = *A*†*Y*.
3. For every cluster of samples, make a crude approximation of a module shared by samples in the cluster. This could be done by, for example, searching for the principal eigenvector of the high-dimensional approximations for each sample in the cluster. Because samples that cluster together are likely to activate similar modules (**fig. S7**), we expect cluster-specific modules to be among the top principal components.
4. Use the modules obtained from each cluster as a crude approximation of the dictionary U.
5. Refine the dictionary, by using a dictionary-learning algorithm with a “local convergence guarantee”. Specifically, we use a variant of SMAF, which we refer to as DL, for “dictionary learning”, that has the property that, if it is initialized “close” to a local optimum, it is mathematically guaranteed to converge to this solution (SOM). Our initial approximations are intended to provide a good starting point for the process. (In some applications (*62*, *63*), there are provable mathematical guarantees that the starting point is good enough to ensure convergence—although we do not yet have such a proof in our application.)
6. Given the refined dictionary, use the methods of compressed sensing to find the sparse activity levels *W* and infer high-dimensional expression values *X*̂.

Interestingly, it has been shown that using different composite measurements for each sample decreases the number of samples needed for the process (*62*).

While the description above closely follows an existing BCS algorithm (*62*), our implementation has modifications that are specifically appropriate for gene expression data. The key difference in our algorithm is that we use SMAF to find the initial sample clusters and modules (SMAF can also be viewed as a bi-clustering algorithm that identifies subsets of genes that co-vary in subsets of tissues). Using SMAF imposes the helpful constraints of dictionary sparsity and non-negativity. The details of this algorithm, which we refer to as BCS-SMAF, can be found in the SOM.

## Blind Compressed Sensing: Results

We tested BCS-SMAF on our 40 datasets. As before, we simulated noisy composite measurements across a randomly selected subset of genes, *Y* = *A*(*X* + *noise*) (*g* = 1000, signal-to-noise ratio: 2), and we varied the number *m* of composite measurements (from 5- to 20-fold fewer than the number of genes). In each case, we compared the gene expression levels recovered from BCS-SMAF with the original values (**Fig. 5B**). (Note that the BCS-SMAF algorithm takes longer to run, and so we decreased the number of genes to run a large number of random simulations in a time efficient manner across all 40 datasets; below we will also discuss BCS-SMAF results with all 14,202 genes in the GTEx dataset.)

Although the performance was expectedly worse than results with a known dictionary, the recovered abundance levels were substantially correlated with the original values. For the GTEx data with *g* = 1000 genes and *m* = 200 composite measurements (thus, 5-fold under-sampling), the expression profiles recovered by BCS-SMAF had an overall Pearson correlation of 82% and Spearman correlation of 66% with the true values, averaged across simulation trials. Increasing the number of measurements had a dramatic effect on recovery performance; for instance, with GTEx, Spearman correlations increased steadily from 40% to 66% with the number of measurements increasing from 25 to 200 (**Fig. 5C**).

We then tested BCS-SMAF on all genes in the GTEx dataset (*g* = 14202; signal-to-noise ratio: 2), again with a number of composite measurements that were 5- to 20 fold fewer than the number of genes. In this case, BCS-SMAF predictions from 5-fold under-sampled (*m* = 2800) data were 78% Spearman correlated with the original values (90% Pearson), and 20-fold under-sampled (*m* = 700) predictions were 59% Spearman correlated (83% Pearson) (**Fig. 5D**).

BCS-SMAF produces good approximations to gene expression, but can we also say that it produces a useful dictionary? Previously, we quantified biological coherence in the module dictionary by calculating the average number of unique gene sets per module. In GTEx data (with all genes, *g* = 14202), SVD and sparse NMF dictionaries had considerably fewer unique gene sets than SMAF (0.15 and 0.06 versus 0.47 unique sets per module). With 5-fold and 20-fold under-sampled BCS-SMAF we find 0.56 and 0.32 gene sets per module, respectively. Thus, we can potentially obtain more insight into the modular structure of gene expression with BCS-SMAF applied to 5- to 20-fold under-sampled data than with the commonly used algorithms of SVD and NMF applied to the entire data.

## Differences between composite measurements and signature gene analysis

We next considered a simple alternative to performing composite measurements consisting of random linear combinations of genes: measuring the levels of individual “signature genes”. In such signature gene analysis (*32*, *33*, *35*, *55*), one selects a small number of individual genes and uses a set of training samples to learn models that can predict the remaining genes from this measured set. Below, we compare the relative advantages of composite measurements vs. signature-gene measurements.

The most obvious advantage of measuring signature genes is conceptual simplicity. Signature-gene measurement is straightforward to understand, and the measurements are relatively simple to implement in practice. However, there is a clear drawback: because the set of signature genes is optimized during model training, the measurement design may change from experiment to experiment as the biological context shifts (for example, if no immune cell types were included in training, but new measurements will be made on these cells, then the signature genes might need to be updated). In contrast, composite measurements can be designed randomly, with a single design that can be suitable for a broad range of contexts.

Generalizing this point, while signature gene methods are restricted to applications in samples that are similar to the samples used to train the model, composite measurements can be used to recover expression levels ‘blindly’ in new samples with BCS-SMAF. Because certain cell types (highly proliferative, transformed cell lines) tend to be more readily available than many other cells (*e.g*., primary, post-mitotic cells), training data may be heavily biased towards the former. It may thus be desirable to avoid relying on training data, especially for studies focused on under-characterized samples.

A common application of signature genes that avoids training bias is pairwise comparison of samples (for example, to identify clusters of samples from a large number of experimental conditions (*64*)). In this case, no imputation is done, and samples are directly compared based only on the measurements of signature genes.

We repeated our earlier analysis to assess how well sample-to-sample distances in high-dimensional space are preserved when calculated from signature genes *vs*. *m* random composite measurements. With *m* = 100, we found that *m* signature genes do not preserve sample-to-sample distances as well as composite measurements (59% vs. 81%) (**fig. S8**).

We next compared the performance of signature genes and composite measurements with respect to their ability to recover unobserved expression levels in testing data, after model building (for signature genes) or dictionary building (for composite measurements) in training data. At moderate levels of noise, the two methods have similar performance. For instance, in GTEx data with 25 measurements and an SNR of 2, the correlation with the original data was 76% with signature genes vs. 82% for composite measurements with a known dictionary (learned from the same data used to choose the signature genes). At higher levels of noise, the performance of signature genes deteriorated more quickly: with an SNR of 0.5, the correlations were 41% vs. 71%. This difference reflects the fact that *composite* measurements tend to cancel out the (uncorrelated) noise in each gene, while signature measurements are highly sensitive to noise in individual genes. In the context of single-cell RNA-Seq, where current methodologies are estimated to sample fewer than 20% of the transcripts in a cell (*39*) (reviewed in (*65*)) this feature is likely to be particularly relevant.

## Making composite measurements in the laboratory: Theory

We now turn to the challenge of implementing compressed sensing in the laboratory—that is, developing a protocol that measures a linear combination of the expression levels of a set of genes, without having to determine the levels of the individual genes. We describe below one approach and present a proof-of-concept experiment, and also address potential alternatives.

Suppose that we want to make the composite measurement consisting of a weighted sum of the expression levels of genes in a set *S*,

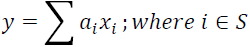

We will initially assume that the weights *a_i_* are all positive.

One approach is to create a pair of a single-stranded DNA probes, *L_i_* and *R_i_*, that hybridize to adjacent locations on the mRNA of gene *i*, so that they may be ligated with an enzyme that selects for RNA-DNA hybrids (such as in (*66*)). The left probe *L_i_* contains a left barcode sequence, and the right probe *R_i_* contains a right barcode sequence that are each the same for all the genes being measured. To perform a composite measurement, one proceeds as follows: (i) create a weighted pool consisting of all of the probe pairs, with their abundance depending on the coefficients *a_i_* in the linear combination; (ii) hybridize the mixture to mRNA in a fixed sample, (iii) perform ligation; (iv) wash away the unhybridized and unligated products; and (v) measure the total amount of the barcode (for example, by qPCR) (**fig. S9A**).

Certain experimental issues must be borne in mind. First, the procedure should be performed so that probe hybridization is linear in the amount of mRNA, and in the concentration of probes. Second, the hybridization efficiency of the probes will not be identical. One should thus “learn” the hybridization efficiencies by performing the procedure on a variety of mRNA samples where the individual gene-expression levels have been determined by a hybridization-independent method, such as RNA-Seq. Moreover, one can use multiple probe pairs per gene, which is likely to be advantageous for single-cell analysis.

While we assumed above that the coefficients were all positive, we can apply the procedure to an arbitrary linear combination by making two separate composite measurements, *y*_+_ and *y*_−_, corresponding to the positive and negative coefficients, and subtract the second from the first.

Finally, we can perform *m* compositional measures simultaneously by hybridizing 2*m* pools corresponding to the positive and negative coefficients of each linear combination and reading out the abundance of the associated barcodes.

We can also imagine performing the final readout by sequencing, rather than by qPCR. In this case we would sequence the final, ligated and purified library of probe pairs, and count the abundance of each barcode. The depth of sequencing required to accurately estimate these relative abundances would then increase as more genes are included in the composite measurements, and as the difference between the most highly expressed and lowly expressed genes grows. On the other hand, with qPCR, the cost is the same whether a measurement spans a single gene, or many thousands. Thus, for composite measures there is an advantage to analog readouts relative to digital alternatives.

Notably, composite measurements can be performed not only for nucleic acids. An alternative approach could leverage methods in mass cytometry (CyTOF) (*67*). CyTOF is similar to traditional flow cytometry, with the modification that antibodies (or *in situ* hybridization probes) are conjugated to heavy metal isotopes rather than fluorophores. Readout is done by time-of-flight mass spectrometry with a fixed number of channels corresponding to ~100 heavy metal ions. Since the number of channels is fundamentally limited, applying compressed sensing to expand the panel of targets by an order of magnitude or more could be transformative for this burgeoning technology.

In order to scale these methods to thousands or tens of thousands of genes, we will need to address practical concerns of building large composite libraries. Our specific choice of random Gaussian measurements follows from analytic, rather than experimental, convenience. Future efforts should focus on developing measurement designs that can be easily and robustly implemented, while also being specifically tailored to constrained reconstruction algorithms. These efforts can be guided by existing work on rational, efficient design of large scale pooled screening (*68*).

## Making composite measurements in the laboratory: Results

We tested this approach by performing a proof-of-concept experiment to measure by a composite approach the levels of 23 transcripts in K562 cells. The genes were selected randomly to capture a large dynamic range of abundance, from highly abundant housekeeping genes (ACTB) to very lowly expressed ones (EMR1). We first evaluated single probes for each gene, using approximately 100 cellular equivalents of RNA. We compared gene expression levels assayed by our protocol (based on ligation, capture and qPCR, for each of the separate probe pairs) with levels assayed by standard qPCR of the mRNA itself. Across three orders of magnitude of dilution for the probes, the correlations varied from 14% to 62% (**table S4**). We then repeated the procedure using four pairs of probes targeting different positions within each gene, all using the same barcode, and observed stronger correlation across the same range of dilutions (ranging from 39% to 88%) (**table S4**). This increase is likely due to robustness conferred by averaging results from different probes with independent noise, and by repeated sampling of a transcript at multiple positions. Finally, we designed 20 arbitrary sets of composite measurements by taking random linear combinations of the 23 genes (**table S5**) and created the corresponding probe libraries. With two replicates of these libraries using different measurement barcodes, we observed 85% and 90% correlation with the expected values calculated directly from the linear combinations of known gene abundances (**fig. S9B**).

We thus find that composite measurements (either across positions in a gene, or across genes) are more robust to noise—demonstrating the potential for improved accuracy with molecular biological compressed sensing. While further tests and optimization will be necessary to extend the methodology to compressed sensing across the transcriptome and to settings such as single-cell analysis, the results here provide an initial proof-of-concept.

## Discussion

Building on established mathematical frameworks, we have demonstrated several perhaps surprising results. The first is that a very small number of randomly designed measurements are sufficient to estimate the similarity between pairs of gene expression profiles. Second, the highly structured nature of gene expression makes it possible to recover a great deal of information – including the expression profiles themselves – from a small number of random composite measurements. Third, we find that an algorithm (SMAF) designed to identify sparse, modular representations of gene expression gives more insight into the underlying biology of a sample than conventional methods of matrix factorization. Finally, without any prior information on the specific modular structure of genes, the mere knowledge that such structure exists is sufficient to blindly extract abundance information from composite observations using a novel algorithm, BCS-SMAF.

We can envision a number of applications of these ideas. For instance, in a pooled genome-wide screen (*e.g.* using CRISPR-Cas9 (*69*)) we might wish to know which genetic perturbations produce a similar response at the level of gene expression (*14*). One way to determine this would be to perform each perturbation in isolation, do RNA-Seq for each condition, and then cluster conditions based on their expression profiles. Our results suggest that making ~100 composite measurements in each condition could be sufficient to produce the same clustering. Alternatively, suppose that composite measurement readout was quantified by fluorescence rather than qPCR (with each measurement corresponding to a different color). We could then perform perturbations in a pool (which is considerably easier that doing a large number in isolation), hybridize probe libraries corresponding to six colors (or however many can be distinguished by FACS), and then sort the cells into a small number of “bins” based on their profiles across the six composite measurements. (This number of measurements would be insufficient to capture whole transcriptome information, but could be informative for a smaller subset of 50-100 genes.) Within each bin we could quantify sgRNA abundance, and perform RNA-Seq in order to associate perturbations of each gene with “characteristic” expression profiles.

Extending this application to recover expression levels directly from composite measurements will be an exciting alternative to RNA-Seq, but there are also a number of other potential use cases for molecular compressed sensing. We mentioned above the possibility of performing readout with CyTOF, which could enable compressive proteomics. This might be particularly interesting in imaging mass cytometry (*70*), in which protein abundance information is spatially resolved to produce an “image” of a fixed sample. With composite measurements, these images could potentially be expanded to include information on thousands of proteins. Moving beyond RNA and protein abundance quantification, we can also consider compressive measurements of other biological systems that might plausibly possess a sparse modular structure. Chromatin landscapes, for example, might be patterned into subsets of the genome that co-vary in chromatin state across a subset of conditions. Other high-dimensional phenotypes such as the spliceosome or metabolome might be similarly structured.

For any of these applications, blind recovery without training data (as in BCS-SMAF) is highly desirable. With our current algorithms, however, there is a considerable, albeit expected, difference in performance in the blind vs. training regime. For example, we were able to achieve ~75% (Spearman) correlation with blind 5-fold under-sampling of GTEx data, and ~90% correlation with 140-fold under-sampling and training data from just 5% of samples. It might therefore be possible to dramatically increase the performance of BCS-SMAF with partial knowledge of gene expression profiles. For example, if we make very weak assumptions on the average expression level of every gene (*e.g.*, approximately highly expressed versus approximately lowly expressed), then we can optimize the measurement designs to be particularly sensitive to lowly-and moderately-expressed genes. Moreover, we can apply features of training data, such as the clustering of genes, to the initialization of BCS without explicitly requiring that the correlation structure remains the same in new samples.

We might also improve these results with several adjustments to the loss function used during optimization. First, for sequencing data we can consider using a Poisson loss, which should be a better model of the process that generated the observed read counts, compared to the squared loss we have used. Second, we can alter the loss function to be specifically sensitive to variations in gene expression across samples (*e.g.*, by optimizing over the average loss for each gene). This would potentially improve the gene-centric correlations, which were lower than other statistics in our results.

We should also find interesting applications and extensions for SMAF, independent of compressed sensing. Here we found that our sparse module dictionaries were more interpretable than those found by SVD or sNMF; this might be particularly useful when rich annotation databases (such as MSigDB (*57*)) are not available to guide our interpretation. We might also extend SMAF to include some notion of a regulatory mechanism, so that the algorithm produces not only a dictionary of modules, but also a set of regulators (such as transcription factors) associated with each module.

Overall we find many intriguing possibilities at the intersection of mathematical theory and molecular biology. To a large extent, biologists are eager to find structure in complex systems that are difficult to access. A large body of mathematical research is about formalizing this problem in the abstract. Biology, therefore, can inform new research directions in mathematics, just as mathematics can inspire new experimental and analytical modalities in biology.

## Acknowledgements

The authors thank Geoffrey Schiebinger and Jonathan Schmid-Burgk for helpful discussions, as well as members of the Regev and Lander labs. B.C. conceived of the project, developed the algorithms, and performed the analysis. L.C. and B.C. designed the experimental implementation, performed the experiments, and analyzed the results. All authors wrote the manuscript. Work was supported by HHMI and the Klarman Cell Obervatory (AR). L.C. was supported by a CRI fellowship.

## List of Supplementary Materials

Supplementary Online Methods (SOM)

Fig S1-S9

References (71-74)

### Additional Files

Tables S1-S5 (Supplemental Tables.xlsx)

